## Supplementary material for "Housing environment bilaterally alters transcriptomic profile in the rat hippocampal CA1 region": S1 Fig

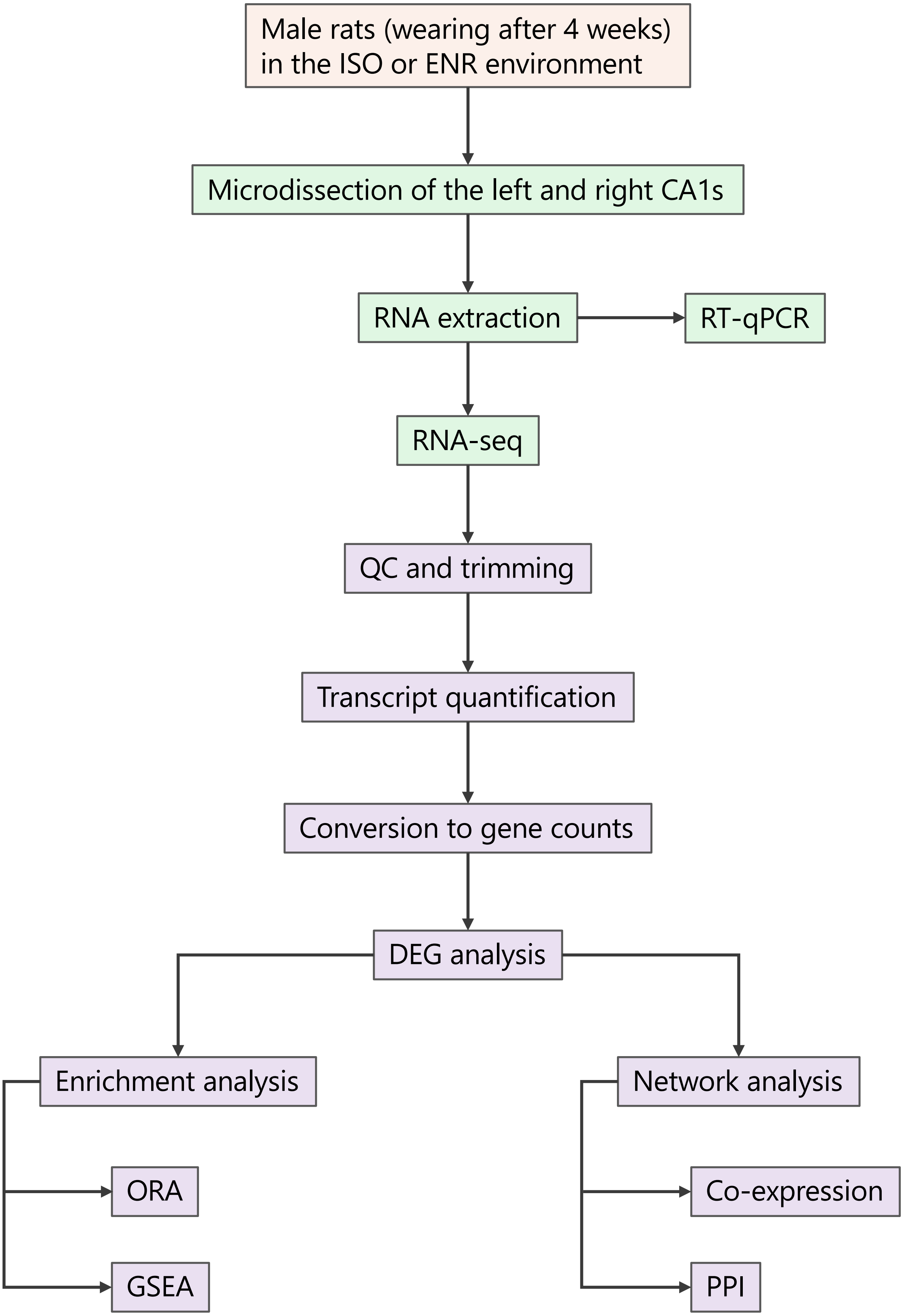


**S1 Fig. Brief workflow chart of this study.**
ISO: isolated. ENR: enriched. DEG: differentially expressed gene analysis. ORA: over-representation analysis. GSEA: gene set enrichment analysis. PPI: protein-protein interaction.
