## Supplementary material for "Housing environment bilaterally alters transcriptomic profile in the rat hippocampal CA1 region": S2 Fig

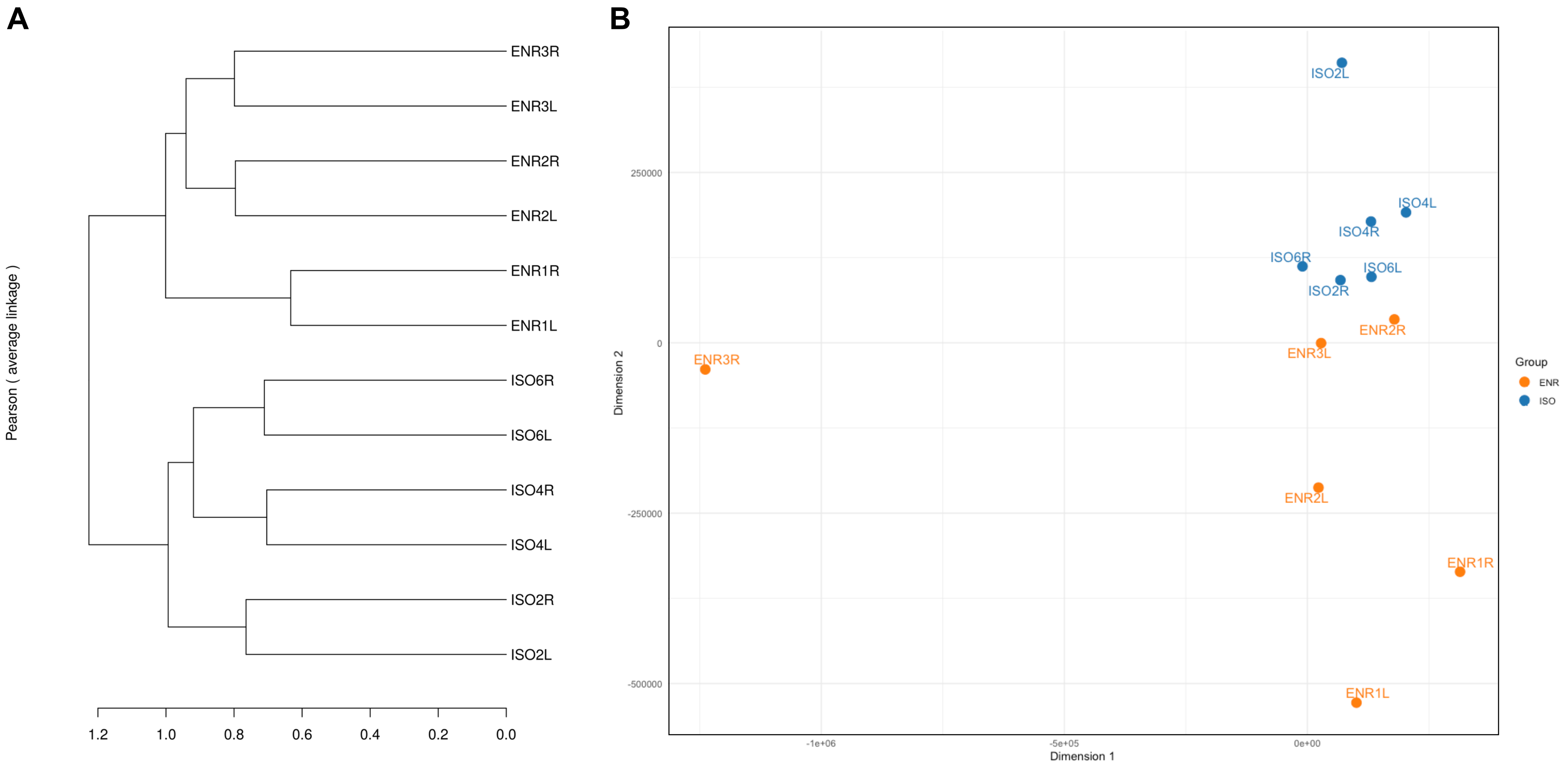


**S2 Fig. Multivariate analyses using the TPM-normalized gene count data.**

**A.** Dendrogram of hierarchical clustering result showing the similarity of the samples using the Pearson's correlation method among the normalized gene count data. ISO: isolated condition, ENR: enriched condition, L: left CA1, R: right CA1. **B.** Multi-dimensional scaling plot visualizing the sample similarities and differences. ENR and ISO groups are colored orange and blue, respectively.
