## Supplementary material for "Housing environment bilaterally alters transcriptomic profile in the rat hippocampal CA1 region": S3 Fig

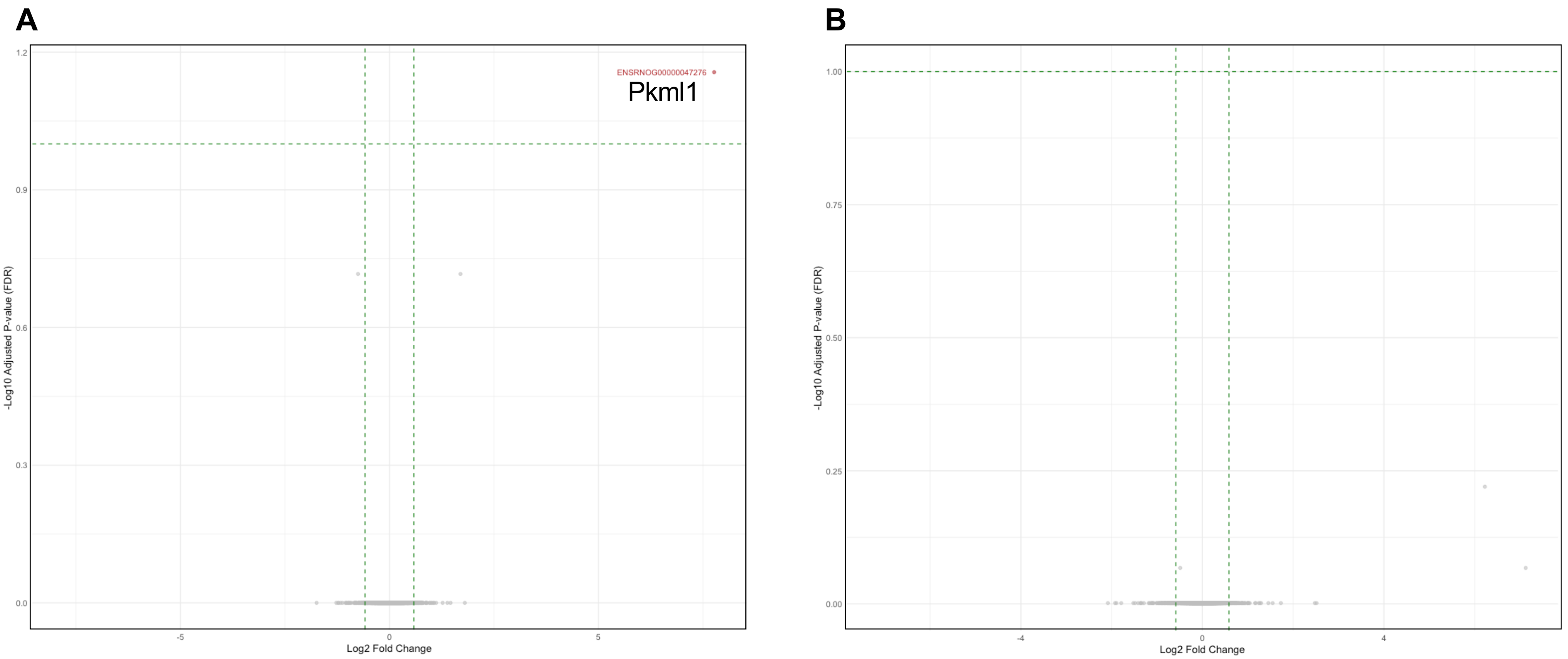


**S3 Fig. Volcano plot visualizations of the left–right comparison.**
FDR: false discovery rate, FC: fold-change. The thresholds ($\log_2$FC = $\pm\log_2 1.5$ and FDR = 0.1) are visualized with green dashed lines. **A.** ISO condition. **B.** ENR condition.
