## Supplementary material for "Housing environment bilaterally alters transcriptomic profile in the rat hippocampal CA1 region": S4 Fig

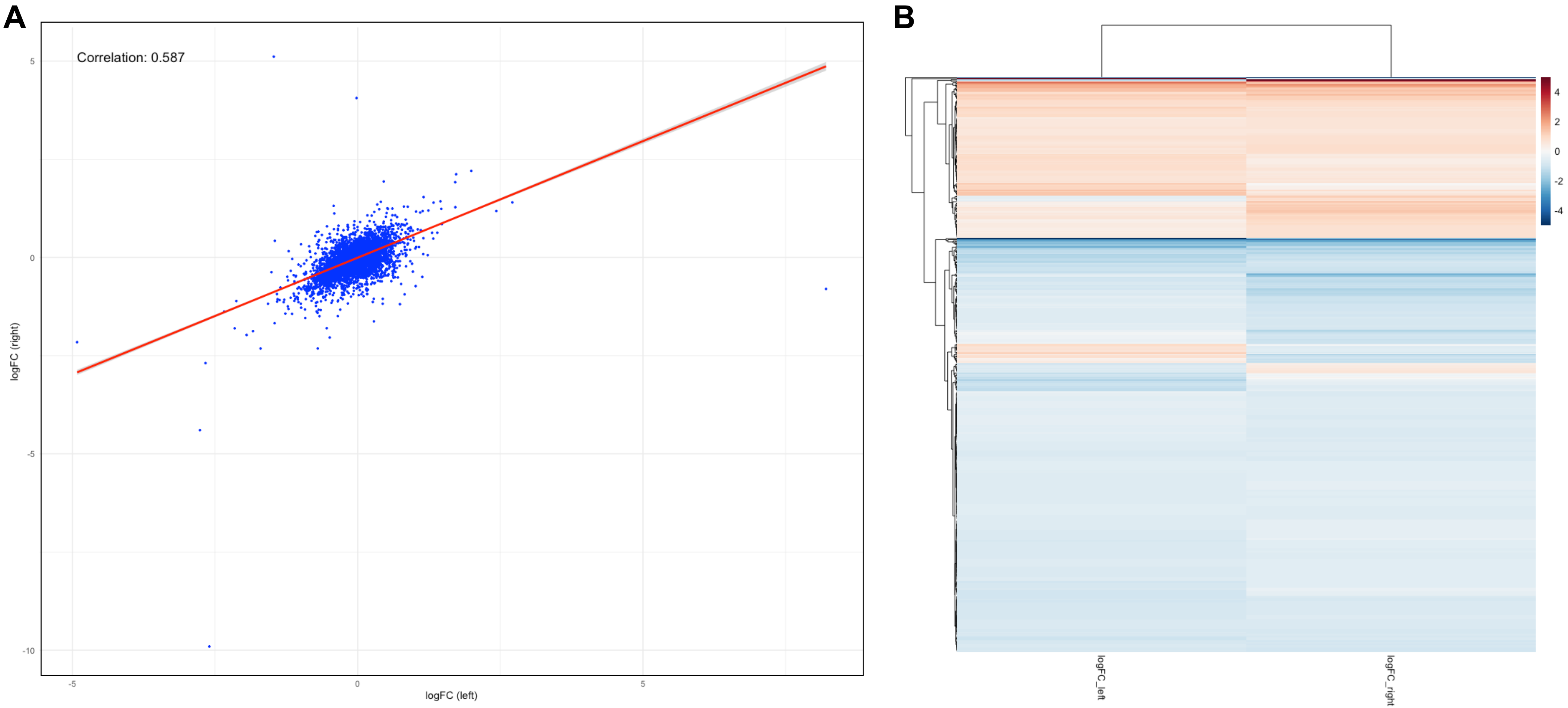


**S4 Fig. Correlation plots focusing on the $\log_2$FC values calculated using edgeR.**
**A.** Scatter plot of the $\log_2$FC values for the environmental comparison in the left and right CA1. **B.** Heatmap of the remarkable $\log_2$FC values for the environmental comparison in the left and right CA1.
