## Supplementary material for "Housing environment bilaterally alters transcriptomic profile in the rat hippocampal CA1 region": S6 Fig

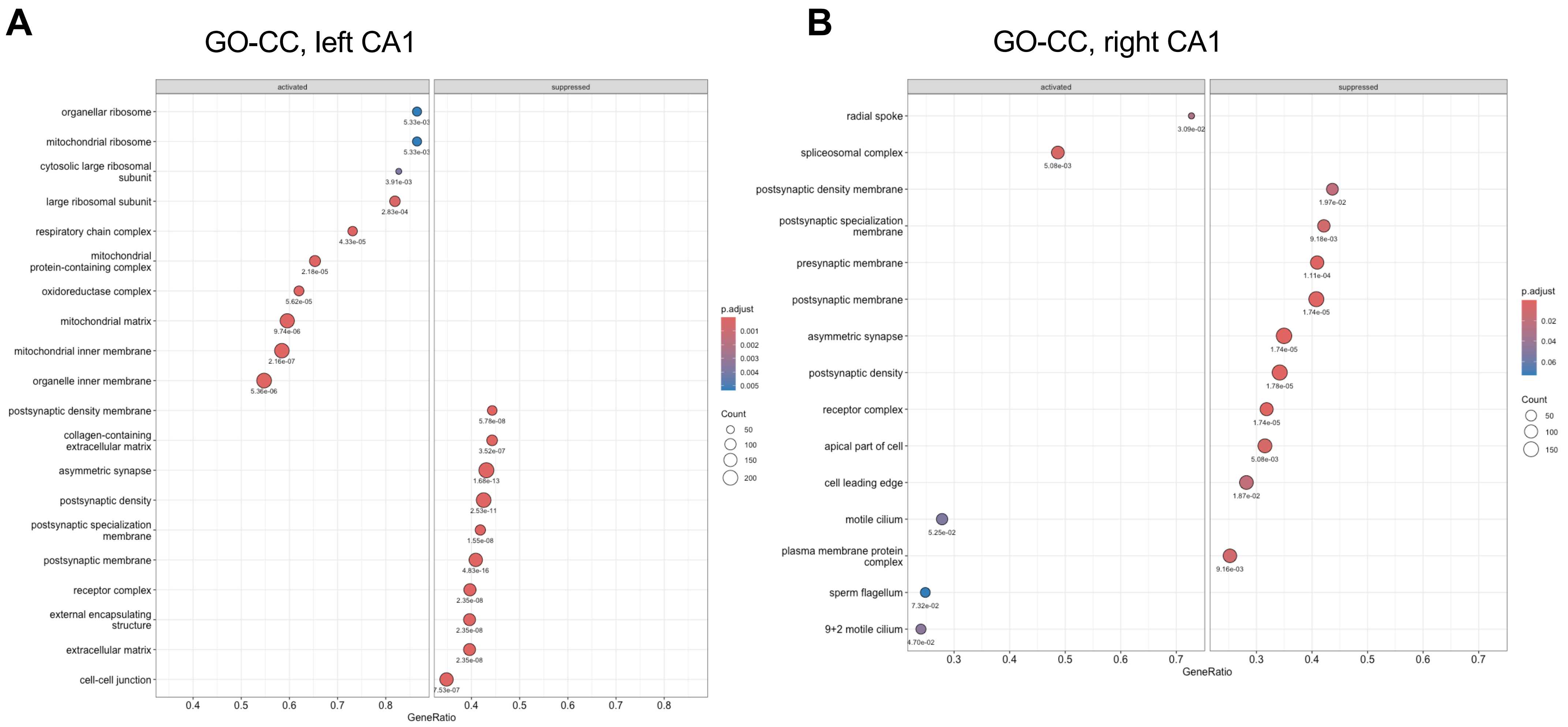


**S6 Fig. GSEA for the environmental comparisons in the left and right CA1 regions using the GO-CC database.**
**A.** Dot plot in the left ISO-ENR. **B.** Dot plot in the right ISO-ENR.
