## Supplementary material for "Housing environment bilaterally alters transcriptomic profile in the rat hippocampal CA1 region": S7 Fig

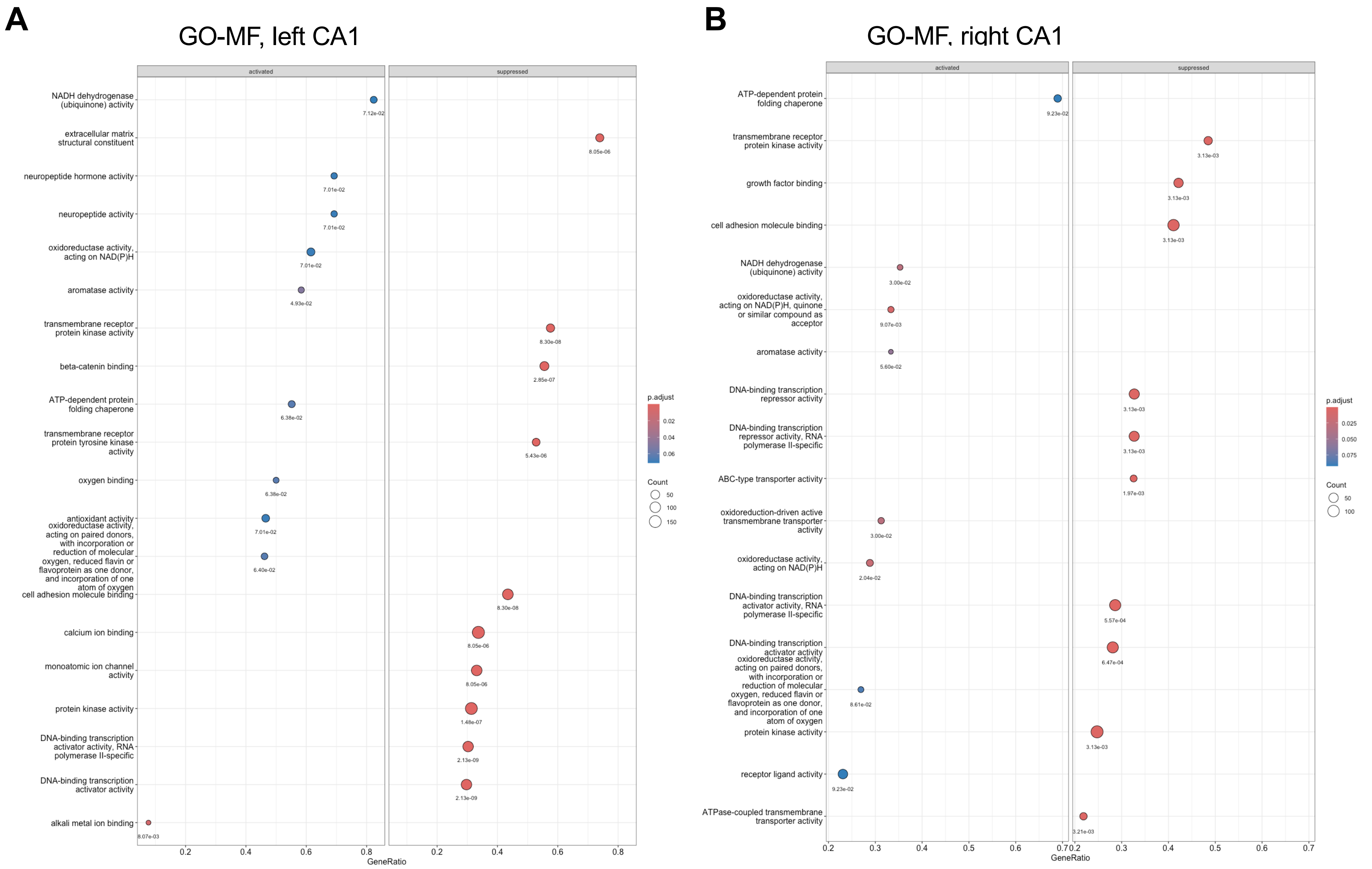


**S7 Fig. GSEA for the environmental comparisons in the left and right CA1 regions using the GO-MF database.**

**A.** Dot plot in the left ISO-ENR. **B.** Dot plot in the right ISO-ENR.
