## Supplementary material for "Housing environment bilaterally alters transcriptomic profile in the rat hippocampal CA1 region": S9 Fig

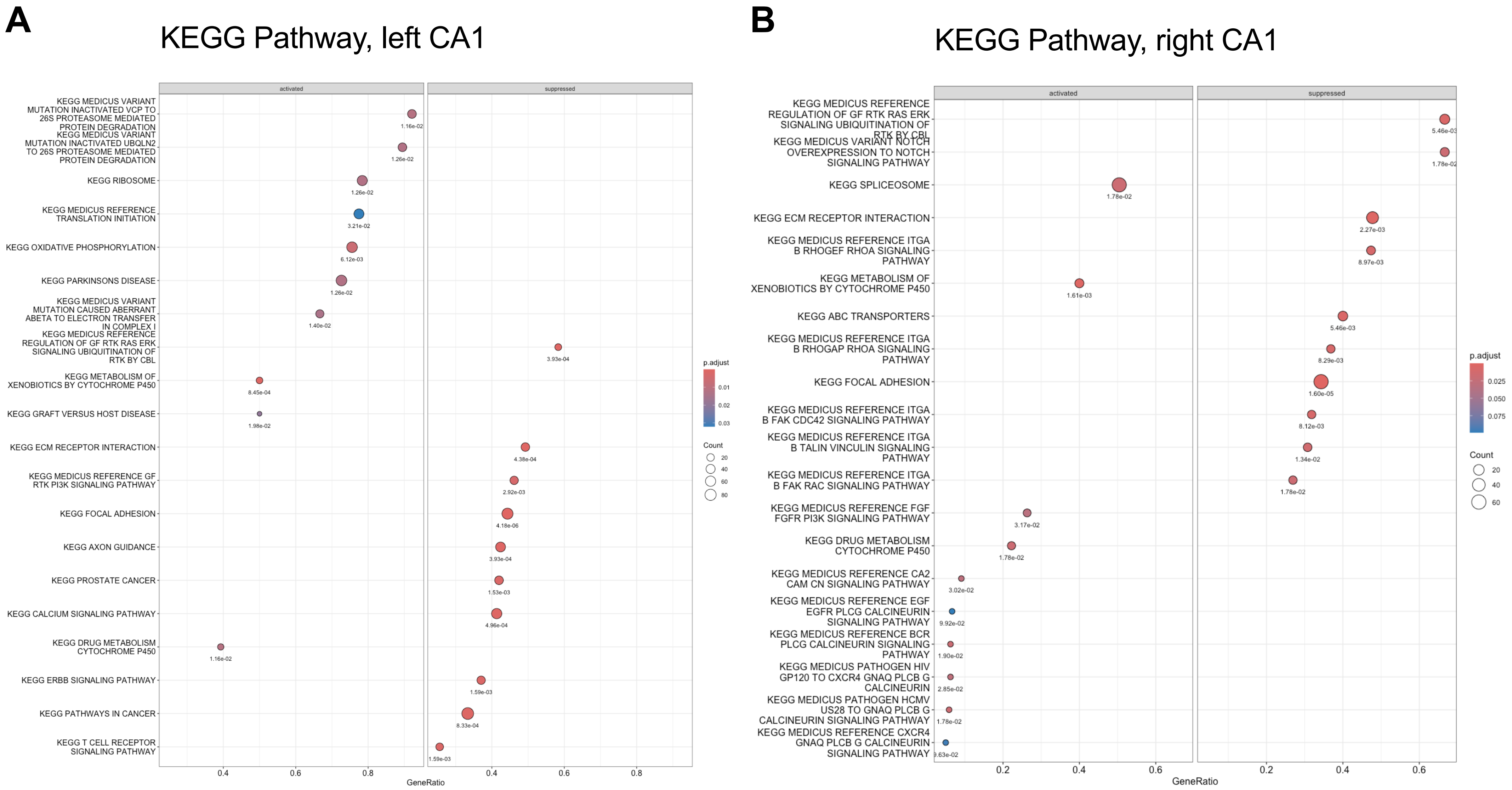


**S9 Fig. GSEA for the environmental comparisons in the left and right CA1 regions using the MSigDB's C2 KEGG Pathway database.**
**A.** Dot plot in the left ISO-ENR. **B.** Dot plot in the right ISO-ENR.
